# QTL Identification for Morphological and Phytochemical Traits in 45 F_2_ Cowpea (*Vigna Unguiculata* (L.) Walp) Mapping Populations

**DOI:** 10.64898/2025.12.16.694557

**Authors:** Henry K. Mensah, Ralieva A.K. Norety, Isaac K. Asante, Felicia Oppong

## Abstract

The research was conducted to study genetic linkage in 45 F_2_ cowpea mapping populations derived from a cross between two inbred lines Golinga (cultivated variety) and a Wild relative. The data used for the study were based on 13 polymorphic SNPs markers, 47 morphological and 23 phytochemical traits (18 amino acids and 5 polyphenols). The morphological variables involved 17 qualitative and 30 quantitative variables of cowpea vegetative and yield and yield-related traits whereas the phytochemical traits involved. Genetic linkage mapping and QTL mapping were performed by using the software QTLiCIMapping version 4.2. All the 17 qualitative markers were polymorphic. Eleven linkage groups were detected at a LOD score of 4. Fourteen markers were distributed on chromosome 1. A total of 79 QTLs were identified which were distributed over four linkage groups. Contributions of the QTLs to percentage phenotypic variation ranged from 0.00% to 27.62% (quercetin). Multiple QTLs located at the same location were identified on chromosomes 1 and 4 indicating potential pleiotropy. QTLs identified for seed related traits were mapped to the same region on chromosome 4. QTLs for the vegetative traits namely, number of branches, number of leaves and branch length were mapped on the same region on chromosome 4. The identification of moderate- to large-effect QTLs on chromosome 4, especially *qSDWG4.1*, offers promising targets for seed weight improvement in cowpea. A total of 42 QTLs were detected for amino acids, most of which were located on chromosome 1, with additional loci on chromosomes 4, 7, and 9. Nine QTLs were identified for phenolic acids. Two QTLs for the flavonoids rutin and quercetin were mapped to a shared region on chromosome 4. This study provided an integrated QTL framework for morphological, reproductive, amino acid, and phenolic traits in cowpea, revealing major genomic regions on chromosomes 1 and 4 that controlled both domesticated-related and nutritional traits. The results offer foundational genomic resources for future fine mapping, candidate gene discovery, and nutritional improvement in cowpea breeding programs.

**Author Summary:** Cowpea is an important food and nutritional security crop, yet many traits that influence its yield, seed quality, and nutrient composition remain poorly understood. In this study, we examined an F₂ population produced by crossing a cultivated cowpea variety (‘Golinga’) with a wild relative to explore genetic factors that control key agronomic and nutritional traits. We measured vegetative, phenological and yield and yield-related traits, 18 amino acids, and five phenolic compounds, and used morphological and SNP markers to identify genomic regions associated with their variation.

We detected 79 QTLs across four chromosomes. Many important seed and reproductive traits were controlled by QTLs on chromosome 4, while chromosome 1 carried major clusters of QTLs for amino acids and phenolic acids. These results reveal genomic “hotspots” that influence both domestication-related traits and nutritional quality.

By identifying the genetic regions associated with these traits, our work provides valuable tools for breeding programs aimed at improving cowpea productivity and enhancing seed nutritional composition. The findings also contribute to a deeper understanding of how wild relatives can introduce beneficial variation into cultivated cowpea.

## Introduction

Genetic-map construction is a critically important tool for further genomic studies, as well as for genetic breeding of economically important species such as cowpea [1]. Cowpea (*Vigna unguiculata* L. Walp) is one of the most ancient crops known to man. Cowpea is also a widely grown legume in tropical and subtropical Africa where it is mostly cultivated by small-scale farmers, usually intercropped with maize or sorghum. The seeds are an excellent source of carbohydrate (50–60%) and an important source of protein (18–35%) [2]. They also contain an appreciable quantity of micronutrients such as vitamin A, iron and calcium [3]. It contains significant amounts of polyphenolic compounds that are beneficial to human and animal health including phenols, flavonoids and tannins [4,5]. Cowpeas contain bioactive antioxidants such as vitamin C, carotenoids and phenolic compounds [4].

Molecular markers are the basis for high-resolution genetic linkage map construction and QTL fine-mapping, which provide powerful tools for genetic analyses of important economic traits such as disease resistance, yield and yield-related traits in cowpea [6]. Linkage maps are estimates of the distance between two genetic loci, based on the frequency of recombination. Highly saturated genetic linkage maps are extremely helpful to breeders and are an essential prerequisite for many biological applications such as the identification of marker-trait associations, mapping QTL, candidate gene identification, development of molecular markers for Marker-Assisted Selection (MAS) and comparative genetic studies [1].

QTL mapping has been reported in most crop plants for diverse traits including yield, seed quality, disease and insect resistance, abiotic stresses like drought and other environmental constraints [7]. Previous studies of grain-related traits in several crops have revealed loci controlling more than one related trait. In Lentil, for example, a flowering time locus was shown to be linked with the grain coat pattern locus [8]. Flowering times influence grain size in different crops [9] Similarly, pre-anthesis changes in vegetative organs can affect the amount of assimilates that are partitioned to the grain during development [7]. Also, post-anthesis processes can affect the time for maturation or grain filling, which could change the grain size [10]. Loci controlling flowering times or other flower morphology traits have also been associated with grain weight or grain size loci in model legume crops [11,12].

The success of QTL mapping depends on the availability and density of the marker maps for the organisms involved [7]. In recent years, comprehensive DNA marker maps have been developed for many crops [13,14]. This facilitates the identification of QTL in crop plants, using an appropriate mapping population. Such populations mostly involve bi-parental progenies such as backcrosses (BC), double haploid lines (DH), F_2_, or recombinant inbred lines (RILs) [15,16]. However, [17] suggested the use of progenies from several parents, to achieve a high probability of obtaining more than one allele at a putative QTL and also to have a more representative estimate of the variance accounted by a QTL. Qualitative characters are common features of natural variation in populations of all eukaryotes, including crop plants. These traits provide a conceptual base for partitioning the total phenotypic variance in terms of additive, dominance and epistatic effects [7]. The basic principle determining whether a QTL is linked to a marker is to partition the mapping population into different genotypic classes based on the genotypes at marker locus and apply relevant statistics to determine whether individuals of one genotype differ significantly from individuals of another genotype [7]. According to [8], the success in using the information on QTLs to increase genetic gain depends greatly on the use of a suitable method to determine the magnitude of QTL effects across multiple environments and whether QTLs are robust across relevant breeding germplasm. Prediction of QTL positions is enhanced by further fine mapping which facilitates testing QTL effects and breeding values in additional populations.

Several types of molecular markers have been developed and advancement in sequencing technologies has geared crop improvement. Genetic mapping uses the Mendelian principles of segregation and recombination to determine the relative proximity of DNA markers along the chromosomes of an organism [1]. The genetic marker is a gene or DNA sequence with a known chromosome location controlling a particular gene or trait. Genetic markers are closely related to the target gene, and they act as assign or flags. Genetic markers are important developments in crop improvement programs such as cowpea.

Thus, this research work aimed at constructing a genetic linkage and QTL maps of cowpea using morphological and SNP in cowpea to identify important yield and phytochemical traits to facilitate the development of improved cowpea varieties to ensure food security and combat malnutrition globally.

## Results and discussion

### Construction of Linkage Maps and Detection of QTLs

The present study identified 79 QTLs in total, 10 QTLs for vegetative traits, 3 QTLs for Phenological, 13 for yield and yield related traits (Table 1) and 53 QTLs for phytochemical traits which involved 18 AAs and 5 polyphenols (Table 2) distributed across 11 linkage groups as shown in Figure 1.Studies such as [19] and [20], [20] similarly mapped their markers to 11 LGs, confirming the stable genome architecture across mapping populations. However, despite these structural similarities, the number, genomic distribution, and effect sizes of QTLs identified in our study differ from earlier reports. [19] used a high-resolution dataset of 34,868 SNPs, mapping 17,996 of them across 11 LGs and detecting 11 QTLs associated with yield-related traits across seven linkage groups. In contrast, [20] employed 215 recombinant inbred lines (RILs) derived from a cross between a cultivated and a wild cowpea accession and evaluated nine domestication-related traits, identifying 16 QTLs across seven chromosomes.

**Figure 1:**
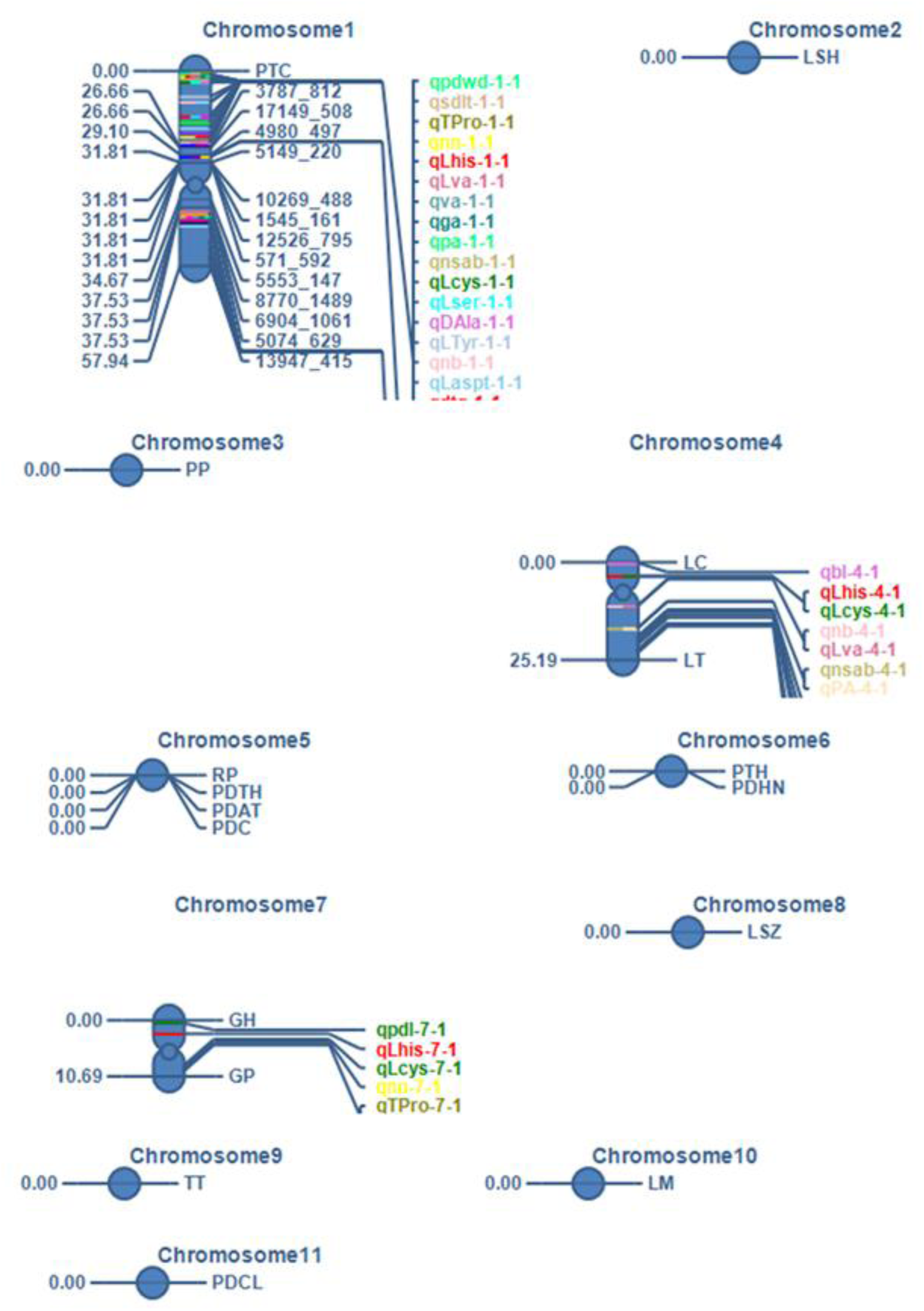
QTLs identified for morphological, amino acids and polyphenols in f2 cowpea mapping populations. Key: GH=Growth habit, GP=Growth pattern, TT=Twinning tendency, PP=Plant pigment, LSZ=Leaf size, LSH=Leaf shape, PLH=Plant hairiness, LC=Leaf colour, LT=Leaf texture, LM=Leaf markings, RP=Raceme position, PDAT=Pod attachment, PDC=Pod curvature, PDTH=Pod thickness, PDCL=Pod colour, PDHN=Pod hairiness, DTG=Mean days to germination, PDL=Mean peduncle length, DFMP=Mean days to first mature pod, NN=Mean number of nodes, NB=Mean number of branches, NL=Mean number of leaves, BL=Mean branch length, DFF=Mean days to first flowering, SPZ=Mean standard petal size,, PDWD= Average pod width, NLPP=Average number of locules per plant, NSAB=Average number of seed aborted, PA=Average percentage seed set, SDLT=Average seed length, SDWG=Seed weight, LHIS=L-Histidine, LCYS=L-Cysteine, LSER=L-Serine, GLU=Glutamine, LLYS= L-Lysine, LASP=L-Asparagine, BTH=B-Threonine, LTH=L-Threonine, LASPT=L-Aspartic Acid, TPRO=Trans-4-Hydroxy-L-Proline, DPRO=D-Proline, LVA=L-Valine, DALA=DL-Alpha-Alanine, LMET=L-Methionine, ILEU=Iso-Leucine, DBLA=DL-Beta_phenyl-Alanin, LTRY=L-Tryptophan, LTYR=L-Tyrosine, VA=Vanillic acid, GA=Gallic acid, PCA=P-coumeric acid, RU=Rutin, QU=Quercetin. QTL name is designated as follow: “q” to indicate quantitative trait loci, followed by the trait name, then followed by the chromosome number, then ‘.’ Finally then the count of traits on a chromosome

**Table 1:**
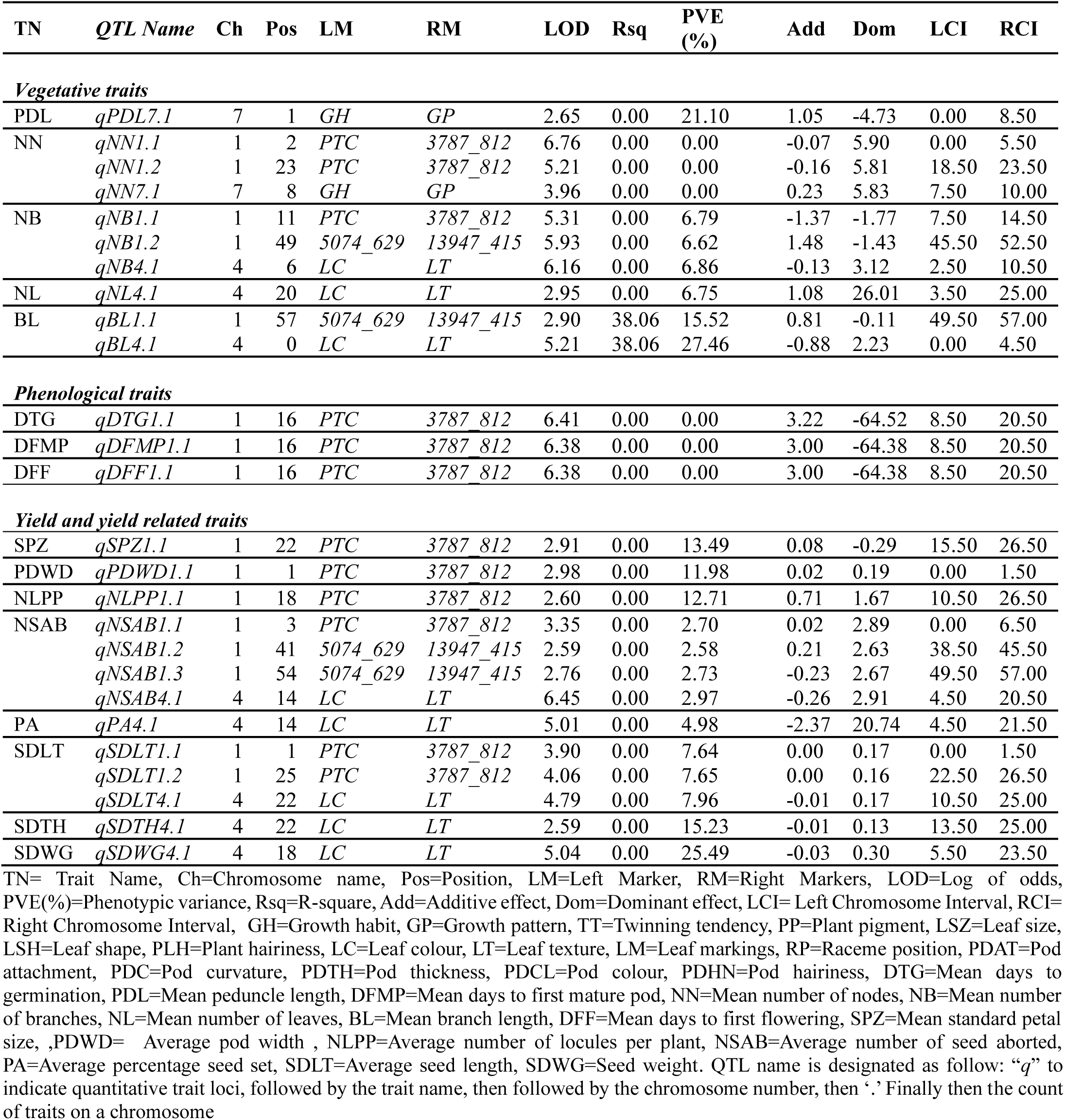
QTLs identified for vegetative, phenological and yield and yield-related morphological traits.

| TN | QTL Name | Ch | Pos | LM | RM | LOD | Rsq | PVE (%) | Add | Dom | LCI | RCI |
| --- | --- | --- | --- | --- | --- | --- | --- | --- | --- | --- | --- | --- |
| **Vegetative traits** |  |  |  |  |  |  |  |  |  |  |  |  |
| PDL | *qPDL7.1* | 7 | 1 | *GH* | *GP* | 2.65 | 0.00 | 21.10 | 1.05 | -4.73 | 0.00 | 8.50 |
| NN | *qNN1.1* | 1 | 2 | *PTC* | *3787_812* | 6.76 | 0.00 | 0.00 | -0.07 | 5.90 | 0.00 | 5.50 |
|  | *qNN1.2* | 1 | 23 | *PTC* | *3787_812* | 5.21 | 0.00 | 0.00 | -0.16 | 5.81 | 18.50 | 23.50 |
|  | *qNN7.1* | 7 | 8 | *GH* | *GP* | 3.96 | 0.00 | 0.00 | 0.23 | 5.83 | 7.50 | 10.00 |
| NB | *qNB1.1* | 1 | 11 | *PTC* | *3787_812* | 5.31 | 0.00 | 6.79 | -1.37 | -1.77 | 7.50 | 14.50 |
|  | *qNB1.2* | 1 | 49 | *5074_629* | *13947_415* | 5.93 | 0.00 | 6.62 | 1.48 | -1.43 | 45.50 | 52.50 |
|  | *qNB4.1* | 4 | 6 | *LC* | *LT* | 6.16 | 0.00 | 6.86 | -0.13 | 3.12 | 2.50 | 10.50 |
| NL | *qNL4.1* | 4 | 20 | *LC* | *LT* | 2.95 | 0.00 | 6.75 | 1.08 | 26.01 | 3.50 | 25.00 |
| BL | *qBL1.1* | 1 | 57 | *5074_629* | *13947_415* | 2.90 | 38.06 | 15.52 | 0.81 | -0.11 | 49.50 | 57.00 |
|  | *qBL4.1* | 4 | 0 | *LC* | *LT* | 5.21 | 38.06 | 27.46 | -0.88 | 2.23 | 0.00 | 4.50 |
| **Phenological traits** |  |  |  |  |  |  |  |  |  |  |  |  |
| DTG | *qDTG1.1* | 1 | 16 | *PTC* | *3787_812* | 6.41 | 0.00 | 0.00 | 3.22 | -64.52 | 8.50 | 20.50 |
| DFMP | *qDFMP1.1* | 1 | 16 | *PTC* | *3787_812* | 6.38 | 0.00 | 0.00 | 3.00 | -64.38 | 8.50 | 20.50 |
| DFF | *qDFF1.1* | 1 | 16 | *PTC* | *3787_812* | 6.38 | 0.00 | 0.00 | 3.00 | -64.38 | 8.50 | 20.50 |
| **Yield and yield related traits** |  |  |  |  |  |  |  |  |  |  |  |  |
| SPZ | *qSPZ1.1* | 1 | 22 | *PTC* | *3787_812* | 2.91 | 0.00 | 13.49 | 0.08 | -0.29 | 15.50 | 26.50 |
| PDWD | *qPDWD1.1* | 1 | 1 | *PTC* | *3787_812* | 2.98 | 0.00 | 11.98 | 0.02 | 0.19 | 0.00 | 1.50 |
| NLPP | *qNLPP1.1* | 1 | 18 | *PTC* | *3787_812* | 2.60 | 0.00 | 12.71 | 0.71 | 1.67 | 10.50 | 26.50 |
| NSAB | *qNSAB1.1* | 1 | 3 | *PTC* | *3787_812* | 3.35 | 0.00 | 2.70 | 0.02 | 2.89 | 0.00 | 6.50 |
|  | *qNSAB1.2* | 1 | 41 | *5074_629* | *13947_415* | 2.59 | 0.00 | 2.58 | 0.21 | 2.63 | 38.50 | 45.50 |
|  | *qNSAB1.3* | 1 | 54 | *5074_629* | *13947_415* | 2.76 | 0.00 | 2.73 | -0.23 | 2.67 | 49.50 | 57.00 |
|  | *qNSAB4.1* | 4 | 14 | *LC* | *LT* | 6.45 | 0.00 | 2.97 | -0.26 | 2.91 | 4.50 | 20.50 |
| PA | *qPA4.1* | 4 | 14 | *LC* | *LT* | 5.01 | 0.00 | 4.98 | -2.37 | 20.74 | 4.50 | 21.50 |
| SDLT | *qSDLT1.1* | 1 | 1 | *PTC* | *3787_812* | 3.90 | 0.00 | 7.64 | 0.00 | 0.17 | 0.00 | 1.50 |
|  | *qSDLT1.2* | 1 | 25 | *PTC* | *3787_812* | 4.06 | 0.00 | 7.65 | 0.00 | 0.16 | 22.50 | 26.50 |
|  | *qSDLT4.1* | 4 | 22 | *LC* | *LT* | 4.79 | 0.00 | 7.96 | -0.01 | 0.17 | 10.50 | 25.00 |
| SDTH | *qSDTH4.1* | 4 | 22 | *LC* | *LT* | 2.59 | 0.00 | 15.23 | -0.01 | 0.13 | 13.50 | 25.00 |
| SDWG | *qSDWG4.1* | 4 | 18 | *LC* | *LT* | 5.04 | 0.00 | 25.49 | -0.03 | 0.30 | 5.50 | 23.50 |

TN= Trait Name, Ch=Chromosome name, Pos=Position, LM=Left Marker, RM=Right Markers, LOD=Log of odds, PVE(%)=Phenotypic variance, Rsq=R-square, Add=Additive effect, Dom=Dominant effect, LCI= Left Chromosome Interval, RCI= Right Chromosome Interval, GH=Growth habit, GP=Growth pattern, TT=Twinning tendency, PP=Plant pigment, LSZ=Leaf size, LSH=Leaf shape, PLH=Plant hairiness, LC=Leaf colour, LT=Leaf texture, LM=Leaf markings, RP=Raceme position, PDAT=Pod attachment, PDC=Pod curvature, PDTH=Pod thickness, PDCL=Pod colour, PDHN=Pod hairiness, DTG=Mean days to germination, PDL=Mean peduncle length, DFMP=Mean days to first mature pod, NN=Mean number of nodes, NB=Mean number of branches, NL=Mean number of leaves, BL=Mean branch length, DFF=Mean days to first flowering, SPZ=Mean standard petal size, PDWD= Average pod width, NLPP=Average number of locules per plant, NSAB=Average number of seed aborted, PA=Average percentage seed set, SDLT=Average seed length, SDWG=Seed weight. QTL name is designated as follow: “q” to indicate quantitative trait loci, followed by the trait name, then followed by the chromosome number, then ‘.’ Finally then the count of traits on a chromosome

**Table 2.** : QTLs identified for amino acids and polyphenolic traits.

| TN | QTL Name | Ch | Pos | LM | RM | LOD | Rsqr | PVE (%) | Add | Dom | LCI | RCI |
| --- | --- | --- | --- | --- | --- | --- | --- | --- | --- | --- | --- | --- |
| **Amino acids** |  |  |  |  |  |  |  |  |  |  |  |  |
| LHIS | *qLHIS1.1* | 1 | 2 | PTC | 3787_812 | 5.67 | 0.00 | 3.94 | -0.01 | 1.37 | 0.00 | 5.50 |
|  | *qLHIS1.2* | 1 | 23 | PTC | 3787_812 | 4.27 | 0.00 | 4.17 | 0.04 | 1.10 | 18.50 | 25.50 |
|  | *qLHIS1.3* | 1 | 52 | 5074_629 | 13947_415 | 2.97 | 0.00 | 4.40 | -0.09 | 1.22 | 47.50 | 52.50 |
|  | *qLHIS4.1* | 4 | 5 | LC | LT | 4.20 | 0.00 | 4.00 | 0.03 | 1.34 | 1.50 | 11.50 |
|  | *qLHIS7.1* | 7 | 5 | GH | GP | 4.67 | 0.00 | 4.09 | 0.05 | 1.30 | 2.50 | 5.50 |
| LCYS | *qLCYS1.1* | 1 | 4 | PTC | 3787_812 | 8.34 | 0.00 | 13.02 | 0.00 | 0.03 | 1.50 | 14.50 |
|  | *qLCYS4.1* | 4 | 5 | LC | LT | 5.70 | 0.00 | 13.01 | 0.00 | 0.03 | 1.50 | 7.50 |
|  | *qLCYS7.1* | 7 | 7 | GH | GP | 5.76 | 0.00 | 13.08 | 0.00 | 0.03 | 3.50 | 10.00 |
| LSEr | *qSER1.1* | 1 | 4 | PTC | 3787_812 | 6.04 | 0.00 | 0.00 | 0.01 | 0.32 | 0.00 | 16.50 |
| GLU | *qGLU1.1* | 1 | 26 | PTC | 3787_812 | 3.83 | 60.51 | 0.00 | 0.01 | -0.36 | 24.50 | 26.50 |
|  | *qGLU1.2* | 1 | 30 | 4980_497 | 5149_220 | 9.62 | 60.51 | 0.00 | 0.01 | 0.60 | 29.50 | 31.50 |
|  | *qGLU1.3* | 1 | 52 | 5074_629 | 13947_415 | 3.66 | 60.51 | 0.00 | -0.01 | 0.21 | 46.50 | 55.50 |
| GLU | *qGLU4.1* | 4 | 22 | LC | LT | 2.65 | 60.51 | 0.00 | -0.02 | 0.25 | 21.50 | 23.50 |
| LLYS | *qLLYS1.1* | 1 | 26 | PTC | 3787_812 | 4.33 | 66.16 | 0.00 | 0.01 | -0.36 | 24.50 | 26.50 |
|  | *qLLYS1.2* | 1 | 30 | 4980_497 | 5149_220 | 10.78 | 66.16 | 0.00 | 0.01 | 0.62 | 29.50 | 31.50 |
|  | *qLLYS1.3* | 1 | 54 | 5074_629 | 13947_415 | 3.07 | 66.16 | 0.00 | -0.02 | 0.21 | 46.50 | 55.50 |
|  | *qLLYS4.1* | 4 | 19 | LC | LT | 4.19 | 66.16 | 0.00 | -0.01 | 0.24 | 18.50 | 23.50 |
| LASP | *qLASP1.1* | 1 | 40 | 5074_629 | 13947_415 | 3.19 | 0.00 | 10.71 | 0.45 | 12.06 | 38.50 | 41.50 |
| BTH | *qBTH1.1* | 1 | 30 | 4980_497 | 5149_220 | 2.78 | 38.51 | 5.70 | 0.01 | 0.13 | 29.50 | 31.50 |
|  | *qBTH1.2* | 1 | 57 | 5074_629 | 13947_415 | 3.39 | 38.51 | 6.01 | -0.03 | -0.04 | 52.50 | 57.00 |
| LTH | *qLTH1.1* | 1 | 49 | 5074_629 | 13947_415 | 4.10 | 0.00 | 14.92 | -0.01 | 0.13 | 41.50 | 54.50 |
| LASPT | *qLASPT1.1* | 1 | 11 | PTC | 3787_812 | 2.97 | 0.00 | 5.17 | -1.40 | -1.75 | 8.50 | 12.50 |
|  | *qLASPT1.2* | 1 | 20 | PTC | 3787_812 | 6.17 | 0.00 | 5.52 | 0.08 | 3.00 | 16.50 | 23.50 |
|  | *qLASPT1.3* | 1 | 44 | 5074_629 | 13947_415 | 5.40 | 0.00 | 5.60 | -0.01 | 2.94 | 40.50 | 47.50 |
| TPRO | *qTPRO1.1* | 1 | 1 | PTC | 3787_812 | 5.67 | 0.00 | 0.00 | 0.03 | 3.49 | 0.00 | 2.50 |
|  | *qTPRO1.7* | 7 | 9 | GH | GP | 4.73 | 0.00 | 0.00 | 0.00 | 3.49 | 7.50 | 10.00 |
| DPRO | *qDPRO1.1* | 1 | 42 | 5074_629 | 13947_415 | 3.03 | 0.00 | 20.89 | 0.06 | 0.52 | 39.50 | 47.50 |
| LVA | *qLVA1.1* | 1 | 2 | PTC | 3787_812 | 3.84 | 0.00 | 2.10 | -0.02 | 0.97 | 0.00 | 4.50 |
|  | *qLVA1.2* | 1 | 25 | PTC | 3787_812 | 3.98 | 0.00 | 2.11 | 0.04 | 0.84 | 21.50 | 26.50 |
|  | *qLVA1.3* | 1 | 39 | 5074_629 | 13947_415 | 3.23 | 0.00 | 2.11 | 0.06 | 0.79 | 37.50 | 41.50 |
|  | *qLVA1.4* | 1 | 55 | 5074_629 | 13947_415 | 3.31 | 0.00 | 2.35 | -0.12 | 0.83 | 52.50 | 57.00 |
|  | *qLVA4.1* | 4 | 6 | LC | LT | 2.54 | 0.00 | 2.15 | -0.06 | 1 | 1.50 | 7.50 |
| DALA | *qDALA1.1* | 1 | 4 | PTC | 3787_812 | 7.65 | 0.00 | 3.15 | -0.01 | 3.42 | 1.50 | 9.50 |
|  | *qDALA1.2* | 1 | 25 | PTC | 3787_812 | 4.55 | 0.00 | 3.15 | 0.06 | 3.42 | 24.50 | 25.50 |
|  | *qDALA1.3* | 1 | 51 | 5074_629 | 13947_415 | 6.92 | 0.00 | 3.13 | -0.04 | 3.39 | 48.50 | 55.50 |
| LMET | *qLMET1.1* | 1 | 41 | 5074_629 | 13947_415 | 3.74 | 0.00 | 3.22 | 0.02 | 0.30 | 38.50 | 45.50 |
|  | *qLMET1.2* | 1 | 55 | 5074_629 | 13947_415 | 4.47 | 0.00 | 3.42 | -0.03 | 0.35 | 50.50 | 57.00 |
|  | *qLMET4.1* | 4 | 17 | LC | LT | 5.37 | 0.00 | 3.26 | -0.03 | 0.36 | 12.50 | 18.50 |
| DBLA | *qDBLA7.1* | 7 | 9 | GH | GP | 15.74 | 0.00 | 0.00 | 0.16 | 4.91 | 5.50 | 10.00 |

**Table 2 : QTLs identified for amino acids and polyphenolic traits**
| TN | QTL<br>Name | Ch | Pos | LM | RM | LOD | Rsqr | PVE<br>(%) | Add | Dom | LCI | RCI |
| --- | --- | --- | --- | --- | --- | --- | --- | --- | --- | --- | --- | --- |
| **Amino acids** |  |  |  |  |  |  |  |  |  |  |  |  |
| LTRY | *qLTRY1.1* | 1 | 43 | 5074_629 | 13947_415 | 3.92 | 0.00 | 26.76 | -0.12 | 16.56 | 42.50 | 47.50 |
|  | *qLTRY9.1* | 9 | 0 | TT | TT | 2.75 | 0.00 | 8.17 | -3.63 | -2.58 | 0.00 | 0.00 |
|  | *qLTYR1.1* | 1 | 9 | PTC | 3787_812 | 7.98 | 0.00 | 17.03 | -0.62 | -0.45 | 4.50 | 14.50 |
| **Polyphenols** |  |  |  |  |  |  |  |  |  |  |  |  |
| VA | *qVA1.1* | 1 | 2 | PTC | 3787_812 | 4.81 | 0.00 | 3.48 | 0.00 | 0.00 | 0.00 | 2.50 |
|  | *qVA1.2* | 1 | 41 | 5074_629 | 13947_415 | 3.08 | 0.00 | 3.24 | 0.00 | 0.00 | 38.50 | 45.50 |
|  | *qVA1.3* | 1 | 55 | 5074_629 | 13947_415 | 2.93 | 0.00 | 3.19 | 0.00 | 0.00 | 49.50 | 57.00 |
| GA | *qGA1.1* | 1 | 2 | PTC | 3787_812 | 4.81 | 0.00 | 3.49 | 0.00 | 0.01 | 0.00 | 2.50 |
|  | *qGA1.2* | 1 | 41 | 5074_629 | 13947_415 | 3.08 | 0.00 | 3.24 | 0.00 | 0.01 | 38.50 | 45.50 |
|  | *qGA1.3* | 1 | 55 | 5074_629 | 13947_415 | 2.93 | 0.00 | 3.19 | 0.00 | 0.01 | 49.50 | 57.00 |
| PCA | *qPCA1.1* | 1 | 2 | PTC | 3787_812 | 4.81 | 0.00 | 3.48 | 0.00 | 0.00 | 0.00 | 2.50 |
|  | *qPCA1.2* | 1 | 41 | 5074_629 | 13947_415 | 3.08 | 0.00 | 3.25 | 0.00 | 0.00 | 38.5 | 45.50 |
|  | *qPCA1.3* | 1 | 55 | 5074_629 | 13947_415 | 2.93 | 0.00 | 3.19 | 0.00 | 0.00 | 49.5 | 57.00 |
| RU | *qRU4.1* | 4 | 22 | LC | LT | 4.33 | 0.00 | 18.08 | 0.00 | 0.02 | 12.5 | 25.00 |
| QU | *qQU4.1* | 4 | 23 | LC | LT | 4.28 | 20.42 | 27.62 | 0.00 | 0.12 | 15.5 | 25.00 |

TN= Trait Name, Ch=Chromosome name, Pos=Position, LM=Left Marker, RM=Right Markers, LOD=Log of odds, PVE(%)=Phenotypic variance, Rsqr=R-square, Add=Additive effect, Dom=Dominant effect, LCI= Left Chromosome Interval, RCI= Right Chromosome Interval, GH=Growth habit, GP=Growth pattern, TT=Twinning tendency, PP=Plant pigment, LSZ=Leaf size, LSH=Leaf shape, PLH=Plant hairiness, LC=Leaf colour, LT=Leaf texture, LM=Leaf markings, RP=Raceme position, PDAT=Pod attachment, PDC=Pod curvature, PDTH=Pod thickness, PDCL=Pod colour, PDHN=Pod hairiness, LHIS=L-Histidine, LCYS=L-Cysteine, LSER=L-Serine, GLU=Glutamine, LLYS= L-Lysine, LASP=L-Asparagine, BTH=B-Threonine, LTH=L-Threonine, LASPT=L-Aspartic Acid, TPRO=Trans-4-Hydroxy-L-Proline, DPRO=D-Proline, LVA=L-Valine, DALA=DL-Alpha-Alanine, LMET=L-Methionine, ILEU=Iso-Leucine, DBLA=DL-Beta-phenyl-Alanin, LTRY=L-Tryptophan, LTYR=L-Tyrosine, VA=Vanillic acid, GA=Gallic acid, PCA=P- coumeric acid, RU=Rutin, QU=Quercetin. QTL name is designated as follow: “q” to indicate quantitative trait loci, followed by the trait name, then followed by the chromosome number, then ‘.’ Finally then the count of traits on a chromosome.

Differences between our results and those reported by [19] and [20] can be attributed to several biological and methodological factors. The population structure is a key determinant: our biparental F₂ population captures only the allelic combinations segregating between two parents, whereas the genetically diverse populations used in the other studies contain far greater recombination and allelic richness. This increases the likelihood of detecting both major- and minor-effect QTLs.

Additionally, variation in statistical thresholds, mapping algorithms, and LOD score criteria across studies contributes to discrepancies in QTL detection. As demonstrated by [21], different interval mapping frameworks vary significantly in power and resolution, leading to study-specific QTL profiles.

### Vegetative traits

#### Peduncle length

A single QTL for peduncle length was detected on chromosome 7, located between markers *GH* and *GP* (0.00–8.5 cM) and peaking at 1.00 cM with a LOD score of 2.65 and an additive effect of 1.05, explaining 21.09% of the phenotypic variation (Table 1, Figure 1)

[20] identified a single QTL *CPedl5* for peduncle length on chromosome 5 that accounted for 71.83% of the variation across a 15.37 cM region in a cultivated × wild cross [20]. The difference in genomic position and effect size between our findings and earlier studies suggests that the chromosome 7 locus detected here may represent a novel allele or population-specific variant absent in previous mapping populations. Such discrepancies across studies highlight the influence of allelic diversity, genetic background, and recombination density on QTL detection, emphasizing the importance of validating peduncle length loci across multiple germplasms sets and environments [20,22,23].

#### Number of branches

Three QTLs associated with number of branches were identified. Two loci, *qNB1.1* and *qNB1.2*, mapped to chromosome 1 within the intervals *PTC* – *3787_812* (7.5 – 14.5 cM) and *5074_629* – *13947_415* (45.5–52.5 cM), peaking at 11.00 cM and 49.00 cM with LOD scores of 5.30 and 5.93, respectively, explaining 6.79% and 6.62% of phenotypic variation. A third locus, *qNB4.3*, mapped to chromosome 4 within *LC – LT* (2.5–10.5 cM), peaking at 6.00 cM with a LOD score of 6.15 and explaining 6.86% of the variation (Table 1, Figure 1). These dispersed loci indicate that multiple genomic regions regulate shoot proliferation in cowpea, consistent with branching QTLs identified in previous cowpea and legume studies [24,25].

#### Number of leaves

One QTL *qNL4.1* for number of leaves was detected on chromosome 4, located between *LC* and *LT* (3.5–25.0 cM) and peaking at 20.00 cM with a LOD score of 2.94 and explaining 6.75% of phenotypic variation (Table 1, Figure 1). Although modest in effect, this locus aligns with earlier findings showing that vegetative growth traits tend to be influenced by several small-effect loci dispersed across linkage groups [23,26].

#### Branch length

Two QTLs for branch length were detected on chromosomes 1 and 4. The QTL *qBL1.1* was found on chromosome 1 and mapped to *5074<u>_</u>629* – *13947<u>_</u>415* (49.5–

57.0 cM), peaking at 57.00 cM with a LOD score of 2.90 and explaining 15.52% of phenotypic variation. The chromosome 4 locus *qBL4.1*, mapped between LC and LT (0.00–4.5 cM), peaked at 0.00 cM with a LOD score of 5.21 and explaining 27.45% of the variation (Table 1, Figure 1). This major-effect region on chromosome 4 aligns with genomic intervals previously associated with shoot elongation and domestication traits in cowpea and related legumes [20,27,28]

Together, these vegetative QTLs spanning peduncle length, leaf production, and branch length reflect both major and minor loci distributed across chromosomes 1, 4, and 7. Their combined effects reinforce the polygenic nature of vegetative architecture in cowpea and highlight genomic regions of interest for marker-assisted breeding aimed at optimizing ease of harvest, canopy structure, plant vigor, and stand establishment.

### Phenological traits (Days to 50% germination, Days to 50% flowering, Days from 50% flowering to mature pods)

A QTL associated with days to 50% germination, *qDTG1.1* was identified on chromosome 1, peaking at 16.00 cM between markers *PTC* and *3787812* (8.5–20.5 cM). This locus had a LOD score of 6.41 but explained 0% of the phenotypic variation. A similar pattern was observed for days to 50% flowering, for which a single QTL, *qDFF1.1* was mapped to the same interval on chromosome 1 with a peak at 16.00 cM, a LOD score of 6.38, and an additive effect of 3.00, though again accounting for 0% of the observed variation. A related QTL associated with days from 50% flowering to mature pods, *qDFMP1.1* was also mapped to this identical region, with a peak at 16.00 cM, a LOD score of 6.38, and 0% variance explained (Table 1, Figure 1)

The co-localization of *qDTG1.1*, *qDFF1.1*, and *qDFMP1.1* suggests the presence of a shared regulatory region influencing early developmental and reproductive timing. However, the negligible phenotypic variance explained by all three loci indicates very weak segregation for these traits in the biparental F₂ population used. This aligns with earlier observations that phenology in cowpea is highly polygenic, governed largely by numerous small-effect loci that are difficult to resolve in low-recombination populations [29].These findings contrast sharply with previous QTL studies employing recombinant inbred lines (RILs), MAGIC populations, and GWAS panels, which revealed multiple flowering-time loci across several chromosomes due to higher allelic richness and more extensive recombination [30,31]. For example, [30], [26] identified flowering-time QTLs on multiple chromosomes, reflecting broader genetic divergence and increased mapping resolution [30]. [20] similarly detected major flowering- and pod-set QTLs on chromosomes 5, 8, and 9 in a cultivated × wild cross characterized by strong allelic contrast [20]. GWAS studies have further shown that flowering time is controlled by numerous small-effect loci dispersed across the genome, many of which escape detection in biparental designs [31].

A direct comparison can be made with the work of [32], who identified eight flowering-time (FT) QTLs in a TV × IT population, including stable multi-environment QTLs on *Vu01* and *Vu08* explaining up to 27.3% of the phenotypic variance [32]. In a separate YA × 58 population, two additional FT QTLs were detected, with multi-environment QTLs on *Vu07* and *Vu09* explaining 13.7–18.4% of trait variation [32]. Similarly, days-to-maturity QTLs (DTM) were identified on *Vu01, Vu02, Vu03, Vu06, Vu08,* and *Vu09.* Compared with these findings, the single weak QTL interval detected in the present study demonstrates that our F₂ population provided limited recombination and insufficient allelic diversity to resolve major phenology loci.

Despite the small effects detected, the consistent alignment of *qDTG1.1*, *qDFF1.1*, and *qDFMP1.1* suggests that this chromosomal region may harbor conserved regulatory genes involved in early development and reproductive transition.

In legumes, flowering transition is regulated through photoperiodic and circadian pathways involving CONSTANS-like (COL) genes, ELF-like transcriptional regulators, and floral integrators such as FT and FLC, many of which map to syntenic or adjacent genomic regions in cowpea and soybean [33–36]

Flowering time is a key adaptive trait influencing drought escape, maturity grouping, and yield stability in cowpea [37,38]. Even small-effect loci such as *qDFF1.1* may hold breeding value when validated in RILs, MAGIC populations, or multi-environment GWAS panels. Future work incorporating these higher-resolution designs will clarify whether this genomic interval contributes environment-specific or epistatic effects affecting early developmental timing.

### Yield and yield-related traits

#### Pod width and Locule Number

One QTL for pod width, *qPDWD1.1* was detected on chromosome 1 between *PTC* and *3787_812* (0.0–1.5 cM), peaking at 1.00 cM with a LOD of 2.97 and explaining 11.98% of the phenotypic variation. Similarly, a QTL for number of locules per pod *qNLPP1.1* mapped to chromosome 1 within the *PTC–3787_812* interval, peaking at 18.00 cM at an of LOD 2.60 and explaining 12.71% of the variation (Table 1, Figure 1). These moderate-effect loci align with findings of [20], who reported pod length and pod structure QTLs distributed across multiple chromosomes, typically explaining 10–25% of trait variance in cultivated × wild RILs [20].

#### Number of seeds aborted per pod and Percentage seed set

Four QTLs influenced number of seeds aborted per pod (NSAB): *qNSAB1.1, qNSAB1.2, qNSAB1.3* on chromosome 1, and *qNSAB4.4* on chromosome 4. These loci had LOD scores between 2.59 and 6.45 but small individual effects (2.58–2.97%) (Table 1, Figure 1). The presence of several low-effect loci suggests strong environmental and polygenic influences on ovule fertility and pod filling. Similar dispersion of small-effect QTLs regulating seed set has been reported in recent multi-environment cowpea mapping studies [32], supporting a model in which reproductive efficiency is controlled by numerous interacting loci rather than a few large-effect genes.

The QTL for percentage seed set, *qPA4.1* also mapped to chromosome 4 flanked on *LC*–*LT* (4.5–21.5 cM), peaking at 14.00 cM of LOD 5.01 with a PVE of 4.97% (Table 1, Figure 1). Genes affecting embryo viability, pollen–pistil compatibility, or pod wall strength may contribute to variation in seed-set success, as shown in comparable QTL studies involving stress-responsive reproductive traits [39,40].

#### Seed length, thickness and weight

In this study, Seed morphological traits were controlled by several QTLs located on chromosomes 1 and 4. For seed length, three QTLs were detected: *qSDLT1.1* at an LOD of 3.90 with PVE of 7.64% and *qSDLT1.2* at an LOD of 4.06 with PVE of 7.64% on chromosome 1, and *qSDLT4.1* with an LOD 4.79 and explained 7.95% of on chromosome 4, indicating polygenic inheritance with small effect sizes. Seed thickness was influenced by a single QTL, *qSDTH4.1* (LOD 2.58; PVE 15.22%) located on chromosome 4. A major QTL for seed weight, *qSDWG4.1* (LOD 5.03; PVE 25.49%), was also found on chromosome 4, accounting for a highest portion of phenotypic variation in yield and yield related traits (Table 1, Figure 1).

The major seed-weight QTL identified in this study, *qSDWG4.1* on chromosome 4, differed from those reported in earlier cowpea studies. [20] detected three high-effect seed-weight QTLs *CSw1, CSw6, and CSw8* and none of their loci mapped to chromosome 4 [20]. In contrast, *qSDWG4.1* did not overlap with any QTL reported in earlier studies, suggesting that the chromosome 4 region detected here may be unique to the wild × cultivated cross used. Other wild × cultivated QTL studies identified seed-weight loci on different genomic regions, with no overlap with [20] or with the QTLs detected in this study [26,41]. These contrasts indicate that seed weight in cowpea is influenced by different sets of loci depending on the genetic background, and that wild germplasm may contribute novel alleles do not present in cultivated varieties.

### Amino Acid (AA) and Polyphenolic Composition

#### Amino Acids

QTL analysis revealed a complex and highly polygenic architecture underlying amino acid (AA) accumulation in the population, with a strong clustering of loci on chromosome 1 and additional contributions from chromosomes 4 and 7, and isolated signals on chromosomes 9. Across all AAs evaluated, a total of forty-two (42) QTLs were detected (Table 2, Figure 1), representing both minor- and moderate-effect loci that collectively shape AA variability.

A prominent pattern was the repeated identification of QTLs on chromosome 1, which contributed to variation in nearly all AAs examined, including L-histidine, L-cysteine, L-serine, glutamine, L-lysine, L-asparagine, β-threonine, L-threonine, L-aspartic acid, D-proline, L-valine, DL-α-alanine, L-methionine, L-tryptophan, and L-tyrosine. Many of these QTLs were localized to overlapping regions flanked by markers *PTC*–*3787_812* and *5074_629*–*13947_415*, indicating the presence of key genomic intervals with pleiotropic or tightly linked genes involved in AA biosynthesis, transport, or metabolism. Notably, several AAs exhibited multiple QTLs within these regions, suggesting either clusters of biosynthetic genes or shared metabolic regulators.

Among the amino-acid traits, L-histidine displayed five QTLs across chromosomes 1, 4, and 7, each explaining approximately 3–4% of the phenotypic variance, indicating a classic minor-effect polygenic pattern. Similarly, glutamine, lysine, and trans-4-hydroxy-L-proline exhibited QTLs with low PVE (≤ 4.00%), reinforcing the quantitative nature of these metabolites. In contrast, several AAs were influenced by moderate-effect QTLs. L-cysteine QTLs on chromosomes 1, 4, and 7 each accounted for ∼13% PVE, marking them as biologically significant loci that may control rate-limiting steps in sulfur-containing AA metabolism. L-asparagine and L-tyrosine also exhibited notable single-QTL effects, with PVE values of 10.7% and 17.0%, respectively.

The QTLs for D-proline and L-tryptophan, *qDPRO1.1* and *qLTRY1.1*, respectively, both mapped to chromosome 1 and exhibited strong effects, with *qDPRO1.1* explaining 20.88% of the phenotypic variance and *qLTRY1.1* accounting for 26.76%, the largest effect observed among all AAs. These results suggest the presence of key loci controlling proline and tryptophan biosynthesis. Although chromosome 1 dominated the QTL contributions, chromosomes 4 and 7 also contributed several meaningful QTLs. Chromosome 4 contained QTLs for L-histidine, L-cysteine, glutamine, lysine, valine, and methionine, many with moderate effects, which highlights this chromosome as a secondary hotspot of AA regulation.

In the present study, three quantitative trait loci (QTLs) associated with L-methionine content—*qLMET1.1*, *qLMET1.2*, and *qLMET4.3* were identified. The first two QTLs, *qLMET1.1* and *qLMET1.2*, were mapped on chromosome 1. These loci exhibited LOD scores of 3.73 and 4.47 and accounted for 3.21% and 3.42% of the phenotypic variance. The third QTL, *qLMET4.3*, was positioned on chromosome 4 between markers *LC* and *LT*, peaking at 17.0 cM, with a LOD score of 5.37, and explained 3.26% of the phenotypic variance. Notably, no QTLs for isoleucine were detected in this biparental F₂ population.

These results contrast with those reported by [42], who utilized a MAGIC (Multi-Parent Advanced Generation Inter-Cross) population comprising 235 F₈ recombinant inbred lines derived from eight genetically diverse founder parents. Their study identified multiple significant QTLs for isoleucine distributed across chromosomes 1, 2, 3, and 6, each with LOD values > 3, collectively explaining up to 9.04% of phenotypic variation. For leucine, QTLs were detected on chromosomes 2, 4, and 5 (LOD > 3; PV = 3.56%). In addition, they reported methionine-associated QTLs on chromosomes 1, 2, 3, and 5, with phenotypic variance contributions reaching 88.43%.

The divergence between our findings and those of may be attributed to several factors, including population structure, allelic diversity, and genetic background effects. The MAGIC population employed in their study captures substantially greater allelic richness, recombination frequency, and epistatic interactions than the biparental F₂ population used here, thereby enabling the detection of a broader range of minor- and major-effect loci. The presence of eight founder parents likely introduced rare alleles and complex gene interactions that are absent in two-parent crosses.

Despite identifying fewer QTLs of relatively small effect, the methionine-associated loci detected in our study particularly those on chromosome 1 partially overlap with regions reported by [42]. This suggests the presence of conserved genomic regions influencing amino-acid metabolism in cowpea and highlights the potential value of integrating diverse population types for future fine-mapping and gene discovery.

Overall, the distribution of QTLs reveals that AA composition in this population is governed by a network of multiple minor-effect loci and several biologically meaningful moderate-effect regions, with chromosome 1 representing the major regulatory hub. The co-localization of QTLs for multiple AAs suggests shared genetic mechanisms, potentially reflecting coordinated regulation of AA biosynthesis pathways. These findings provide strong candidate genomic regions for downstream marker validation, metabolic pathway analysis, and breeding strategies aimed at improving AA profiles.

#### Polyphenols

QTL mapping for phenolic compounds revealed strong evidence of trait clustering on chromosome 1 for simple phenolic acids and a distinct concentration of high-effect QTLs on chromosome 4 for flavonoids. In total, eleven QTLs were detected across four phenolic metabolites and represented both minor-effect loci associated with phenolic acids and major-effect loci governing flavonoid accumulation (Table 2, Figure 1).

Three structurally related phenolic acids exhibited nearly identical QTL patterns as for follows vanillic acid (*qVA1.1*, *qVA1.2, qVA1.3)*, gallic acid (*qGA1.1*, *qGA1.2, qGA1.3*), and p-coumaric acid (*qPCA1.1*, *qPCA1.2, qPCA1.3*) ). QTLs involved with each compound were all mapped to chromosome 1, and all localized to similar marker intervals. These QTLs consistently occurred within two major genomic regions: a proximal region near *PTC*–*3787*_812 (0–2.5 cM) and two distal regions within *5074*_*629*–*13947*_*415*, spanning 38.5–57.0 cM. The repeated detection of QTLs within these intervals suggests the presence of shared regulatory loci or tightly linked genes governing early-pathway phenolic acid biosynthesis. Individual QTL effects were small, explaining 3.19–3.48% of phenotypic variation for each compound, consistent with the minor-effect, quantitative nature of phenolic acid accumulation. The near-identical genomic positions and PVE values across all three acids further indicate coordinated genetic control of these core metabolites.

In contrast to the simple phenolic acids, the flavonoids rutin and quercetin displayed a distinctly different genetic architecture. Each trait was governed by a single, moderate-to-major effect QTL, both mapped to chromosome 4 in a shared genomic interval flanked by *LC*–*LT.* The QTL for rutin, *qRU4* peaked at 22.0 cM and explained 18.08% of phenotypic variation, while the QTL for quercetin, *qQU4* peaked at 23.0 cM and accounted for an 27.61% of phenotypic variance. These high-effect QTLs likely represent key regulatory loci associated with branch-point genes in the flavonoid pathway, such as those involved in glycosylation or flavonol conversion. The tight proximity of the rutin and quercetin QTLs strongly suggests either pleiotropy or the presence of linked biosynthetic genes, reflecting the biochemical relationship between the two flavonoids.

Together, these results reveal a strong division in the genetic control of phenolic compounds: chromosome 1 serves as the primary hub for simple phenolic acids, while chromosome 4 contains major regulatory regions for flavonoids. This pattern mirrors known biochemical pathway structure, in which upstream shikimate-derived phenolic precursors are broadly regulated, while downstream flavonoid products are often controlled by specific high-impact loci. The identified QTL regions therefore represent powerful genomic targets for metabolic improvement and for dissecting the regulatory networks that differentiate core phenolic acids from specialized flavonoids.

### Comparative Interpretation Across Studies and Breeding Implications

Chromosome 4 consistently carries large-effect QTLs for seed size and weight, matching domestication-related findings in both RIL and MAGIC populations [20–23]. Effect sizes in this biparental F₂ population are moderate, as expected given fewer recombination events compared with RIL/GWAS/MAGIC populations, which typically reveal more QTLs with clearer resolution [23]. Environmental sensitivity influences reproductive traits, such as seed abortion and seed set, which often show low variance explained yet multiple contributing loci [43].

The identification of moderate- to large-effect QTLs on chromosome 4, especially *qSDWG4.1*, offers promising targets for seed weight improvement in cowpea. Meanwhile, the smaller-effect QTLs on chromosome 1 highlight opportunities for selection aimed at reducing seed abortion and improving reproductive efficiency under stress.

The distribution of amino acid QTLs particularly methionine and valine mirrored patterns reported in multi-parent studies. Compared with the broad, high-variance QTL networks detected in the MAGIC population of [42], the present biparental study identified fewer methionine QTLs of moderate effect on chromosomes 1 and 4. However, partial overlap on chromosome 1 suggests conserved genomic regions influencing amino acid metabolism across population types. Differences in QTL number and effect size reflect contrasting allelic diversity and recombination structure between two-parent F₂ and eight-parent MAGIC populations.

Polyphenolic QTLs showed similar partitioning, with vanillic, gallic, and p-coumaric acids mapping to chromosome 1 and flavonoids such as rutin and quercetin mapping to chromosome 4, reinforcing these chromosomes as key regulatory hubs for both morphological and biochemical traits.

The consistent identification of moderate- to large-effect QTLs on chromosome 4 particularly *qSDWG4.1* for seed weight and *qLMET4.3* for methionine provides valuable targets for marker-assisted selection. Chromosome 1, with its dense clusters of reproductive, amino acid, and polyphenolic QTLs, offers additional opportunities for improving seed quality and stress resilience. Multi-environment validation and genomic selection approaches will be essential for capturing the polygenic basis of nutritional traits and ensuring stability across diverse production environments.

## Materials and Methods

### Development of mapping population and Phenotyping

Forty - five F_2_ mapping populations (accR1-accR45) of cowpea were developed from a biparental cross between Golinga (accR46) and a Wild relative (accR47). These accessions were obtained from the Genetics Lab in the Department of Plant and Environmental Biology (DPEB). A single row plot design was used. Two seeds from each F_2_ population and the two parental checks (Golinga and Wild) were planted in a pot. A distance of 1m x 1m was maintained between the pots. The 5-row plots consisted of 10 pots.

Phenotypical data collected were grouped under morphological and phytochemical traits as presented in **Table 3**. Morphological characterization was conducted under the two broad areas qualitative and quantitative traits. The standard Descriptor for Cowpea by the International Board for Plant Genetic Resource [44] was used. Eighteen morphological qualitative traits were scored based on 11 vegetative and 7 reproductive traits. Twenty-nine (29) morphological quantitative traits measured were grouped based on 10 vegetative, 3 phenological and 16 yield and yield-related (reproductive traits). Twenty-three phytochemical traits were measured, which involved 18 amino acid and 5 polyphenols.

**Table 3:** Phenotypic traits evaluated for the construction of linkage maps and detection of QTLs.

| Phenotypic traits | Group | Trait | Symbol |
| --- | --- | --- | --- |
| **Morphological qualitative traits/markers** | *Vegetative traits* | Growth habit | *GH* |
|  |  | Growth pattern | *GP* |
|  |  | Twinning tendency | *TT* |
|  |  | Plant pigmentation | *PP* |
|  |  | Leaf size | *LSZ* |
|  |  | Leaf shape | *LSH* |
|  |  | Plant hairiness | *PLH* |
|  |  | Leaf colour | *LC* |
|  |  | Leaf texture | *LT* |
|  |  | Leaf markings | *LM* |
|  |  | Petiole thickness | *PTH* |
|  | *Yield and yield-related traits* | Raceme position | *RP* |
|  |  | Pod attachment | *PDAT* |
|  |  | Pod curvature | *PDC* |
|  |  | Pod thickness | *PDTH* |
|  |  | Pod colour | *PDCL* |
|  |  | Pod hairiness | *PDHN* |
|  |  | Pod tip colour | *PTC* |
| **Morphological quantitative traits** | *Vegetative traits* | Plant height | *PH* |
|  |  | Number of nodes | *NN* |
|  |  | Leaf length | *LL* |
|  |  | Leaf width | *LW* |
|  |  | Number of leaves | *NL* |
|  |  | Stipule length | *STL* |
|  |  | Stipule width | *STW* |
|  |  | Peduncle length | *PDL* |
|  |  | Number of branches | *NB* |
|  |  | Petiole length | *PTL* |
|  | *Phenological traits* | Days to germination | *DTG* |
|  |  | Days to first flowering | *DFF* |
|  |  | Days to first matured pod | *DFMP* |
|  | *Yield and yield-related traits* | Standard petal size | *SPZ* |
|  |  | Keel/Lobe length | *LBL* |
|  |  | Number of racemes per plant | *NRPP* |
|  |  | Number of pods per peduncle | *NPPPD* |
|  |  | Number of pods per plant | *NPPPL* |
|  |  | Pod length | *PDLN* |
|  |  | Pod width | *PDWD* |
|  |  | Number of locules per pod | *NLPP* |
|  |  | Number of seeds per pod | *NSPP* |
|  |  | Number of seeds per plant | *NSPPL* |
|  |  | Number of seed aborted | *NSAB* |
|  |  | Percentage seed set | *PA* |
|  |  | Seed length | *SDL* |
|  |  | Seed width | *SDWT* |
|  |  | Seed thickness | *SDTH* |
|  |  | Seed weight | *SDWG* |

**Table 3: Phenotypic traits evaluated for the construction of linkage maps and detection of QTLs (cont'd)**
| Phenotypic traits | Group | Trait | Symbol |
| --- | --- | --- | --- |
| Phytochemical traits | *Amino Acid* | L-Histidine | *LHIS* |
|  |  | L-Lysine | *LLYS* |
|  |  | L-Cysteine | *LCYS* |
|  |  | L-Methionine | *LMET* |
|  |  | L-Aspartic acid | *LASPT* |
|  |  | L-Valine | *LVA* |
|  |  | L-Threonine | *LTH* |
|  |  | L-Serine | *LSER* |
|  |  | L-Tryptophan | *LTRY* |
|  |  | L-Tyrosine | *LTYR* |
|  |  | L-Asparagine | *LASP* |
|  |  | D-Proline | *DPRO* |
|  |  | Glutamine | *GLU* |
|  |  | Isoleucine | *ILEU* |
|  |  | DL-Beta-phenylalanine | *DBLA* |
|  |  | DL-Alpha-alanine | *DALA* |
|  |  | Trans-4-hydroxyl-proline | *TPRO* |
|  |  | B-Threonine | *BTH* |
|  | *Polyphenols* | Gallic acid | *GA* |
|  |  | Vanillic acid | *VA* |
|  |  | p-Coumaric acid | *PCA* |
|  |  | Rutin | *RU* |
|  |  | Quercetin | *QU* |

#### Amino Acid (AA) Analysis

##### Sample preparation

Standard solution was prepared by weighing 2.5 mg of the standard compound into 25 ml of ethanol and diluted to a concentration of 100 ppm. A mixed standard solution of 10 ppm was prepared from the stock solution which was diluted to 1ppm as the working solution. A weight of 0.5 g of cowpea flour was extracted in 20 ml of 100 % methanol.

##### Chromatography

Column used was Agilent Poroshell 120 bonus RP 2.7 um, 2.1 x 100 mm; flow rate was 0.15 ml/min. Mobile phases were: 100 % water, 0.1 % formic acid (A) and 100 % methanol, 0.1 % formic acid (B). The gradient was 0 to 5 % (B) over 5 minutes and 5 to 50 % (B) over 15 minutes. The injection volume was 6 μl.

##### Mass spectrometry

MS-Aligent Tripple quadrupole 6420 was used to identify and quantify.

#### Polyphenols

##### Preparation of sodium carbonate concentration

A 50 ml volumetric flask was filled with 20 ml distilled water. A mass of 6.25 g of sodium carbonate was weighed and dissolved in the distilled water. The solution was boiled, allowed to cool and then a few crystals of sodium carbonate was added. The solution was made to stand for 24 hours and then filtered. Distilled water was added to reach the 25 ml mark.

##### Extraction of samples for phytochemical studies

About 20 cowpea seeds from pods of each of the plants harvested were ground into powder using the mortar and pestle. A mass between 0.25g and 0.5g of the flour was weighed poured into McCartney bottles. Between 10ml and 20 ml of 100% ethanol was added and shaken. The bottles were covered with aluminium foil and allowed to stand for 24 hours. After 24 hours, the sample in solution was filtered and the filtrate was stored in tightly covered McCartney bottles at a temperature of 4°C in the fridge.

##### Determination of phenolic acid content

Extract of the cowpea flour for each plant harvested was analysed for phenolic compounds using the Folin-Ciocalteu method [45].. Two (2) millilitres of each extract were measured with a measuring cylinder and then diluted to 20 ml with distilled water in test tubes. Twenty microliters (20μl) of diluted samples were pipetted into cuvettes. A volume of 1.58 ml of distilled water and 100 μl Folin-Ciocalteu reagent was measured with a measuring cylinder and a 100 μl micropipette respectively and added to the solution. The solution was shaken to mix. A volume of 300 μl of sodium carbonate was pipetted with a micropipette and added to the solution after about 5 minutes and shaken. The solution was placed in an oven for 30 minutes at a temperature of 40°C. The cuvettes were taken out after 30 minutes and allowed to stand until they cooled. The absorbance at 765 nm was determined against the blank ethanol using the visible spectrophotometer. The concentration for the phenolic compounds was determined from the standard curves of linear equations.

##### Determination of flavonoid content

The modified aluminium chloride calorimetric procedure was used for the determination of flavonoid content in the cowpea samples. A volume of 100 μl of sample extract was pipetted and added to 500 μl of distilled water and 30 μl of 5% sodium nitrite in cuvettes. The resulting solution was made to stand for about 5 minutes after which 30 μl of 10% aluminium chloride was added. The solution was allowed to stand again for 6 minutes after which 200 μl of 1M sodium hydroxide and 110 μl of distilled water were added and vortexed. Measurement of absorbance of the solution was made at a wavelength of 425nm and 415nm for both rutin and quercetin using the spectrophotometer. The concentration for individual flavonoid compounds was calculated according to their respective standard curves and the results expressed as mg/l of extract.

##### Standard curves

The standard curves generated for the quantification of the polyphenolic compounds demonstrated strong linear relationships between concentration and absorbance (R² ≥ 0.90). The phenolic standards gallic acid, vanillic acid, and p coumaric acid yielded the calibration equations **y = 0.2263x + 0.1606**, **y = 0.1246x + 0.0794**, and **y = 0.0726x + 0.0874**, respectively. The flavonoid standards rutin and quercetin produced the equations **y = 0.0229x + 0.0250** and **y = 0.1548x + 0.0301**, respectively. These calibration curves formed the analytical basis for determining the concentrations of the corresponding polyphenolic compounds in the cowpea extracts.

### Genotyping

#### Genotyping with single nucleotide polymorphism (SNPs) markers

Four leaf discs of 6 mm diameter were punched from 15-day old leaves and collected into 96 deep well PCR plates. Before sampling, leaves were dried at 50°C for 2 days to prevent possible mould development during shipment to the outsourced laboratory. DNA extraction and genotyping were done by sequencing in the United Kingdom hosted by the Laboratory of the Government Chemist, LGC. LGC genomics used a unique proprietary in-house technology (oKtopure™ protocol) to extract the total DNA. Isolated DNA was analyzed using UV spectrophotometry to estimate both the quality and quantity of the DNA, while preliminary PCR at a serial dilution was carried out and the results produced were duplicated to identify the best dilution to run the samples. The SNP genotyping was done using KASP genotyping reactions. KASP™ is also proprietary genotyping technology of LGC TM. It consists of three components namely the sample DNA, KASP assay mix and KASP master mix. KASP master mix contains two universal (FRET) fluorescent resonance energy transfer cassettes (FAM and HEX), ROX™ passive reference dye, Taq polymerase, free nucleotides and MgCl2 in an optimized buffer solution, while the KASP assay mix is specific to the targeted SNP and consists of two competitive, allele-specific forward primers and one common reverse primer. Each forward primer incorporates an additional tail sequence that corresponds to one of two universal FRET cassettes present in the KASP master mix.

#### Construction of Linkage Maps and Detection of QTLs

Screening was done to select polymorphic morphological traits and SNPs markers within the mapping populations. Out of 43 SNPs markers that were screened thirteen showed polymorphisms (Table 4). Eighteen polymorphic morphological markers were scored (Table 3). The QTL IcMapping software version 4.2 [14] was used for the linkage map construction. The ‘Group’ command was used to identify linkage grouped and the ‘Order’ command was used to establish the most likely order within each linkage group. The orders were confirmed by permuting all adjacent markers by the ‘Ripple’ command. LOD score of 4.0 was used as the threshold for detection of linkage. The analysis was performed by using the Inclusive Composite Mapping with additive and dominance ICIM ADD method.

**Table 4:**
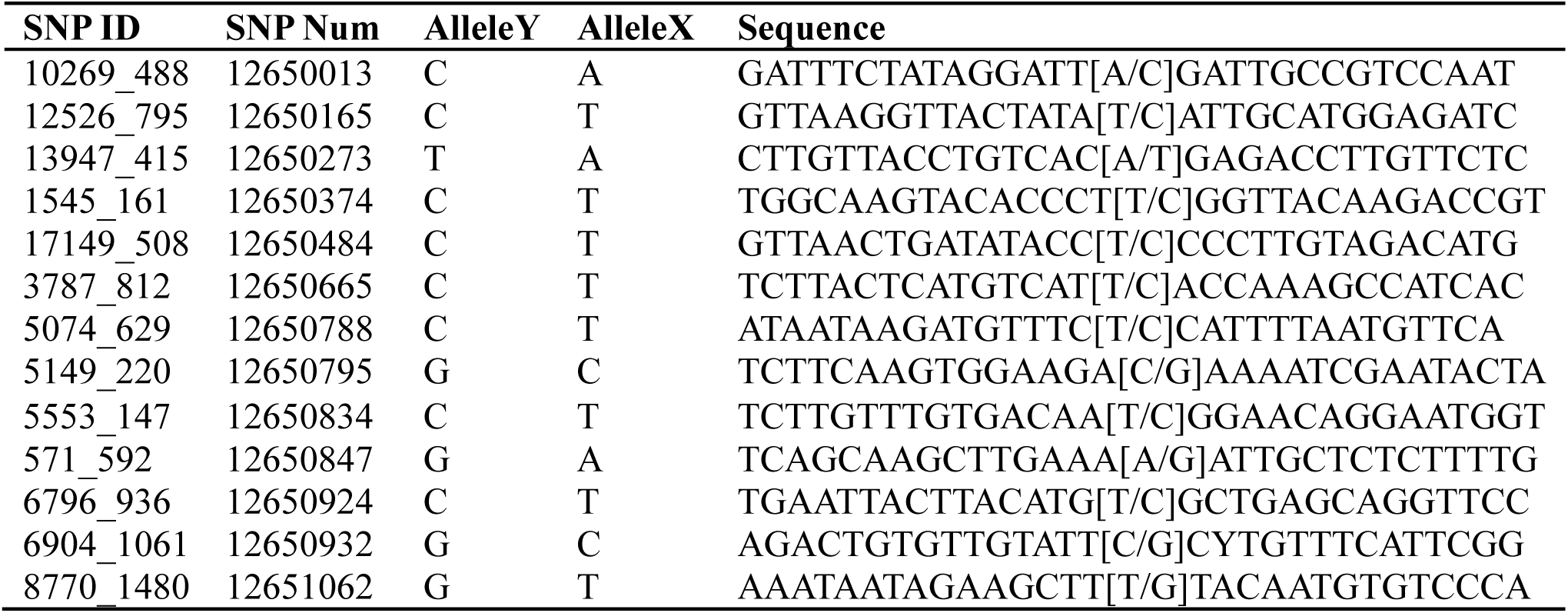
SNPs markers used for Linkage map construction.

The QTL IcMapping software version 4.2 [47] was used for the linkage map construction. The ‘Group’ command was used to identify linkage grouped and the ‘Order’ command was used to establish the most likely order within each linkage group. The orders were confirmed by permuting all adjacent markers by the ‘Ripple’ command. LOD score of 4.0 was used as the threshold for detection of linkage. The analysis was performed by using the Inclusive Composite Mapping with additive and dominance ICIM ADD method. A threshold LOD score of 4.0 used to declare significant QTL.

Quantitative trait loci map construction was performed by using the QTL IcMapping software version 4.2. Quantitative traits were made up of 31 morphological quantitative traits, eighteen amino acid compounds and five polyphenolic compounds (Table 3). QTL map was constructed using the eighteen morphological markers and 13 SNPs markers. The morphological qualitative data and SNP data were used to construct a genetic linkage map for the F_2_ mapping population of cowpea derived from biparental cross Golinga and Wild relatives. Only traits for which the progenies showed significant differences were maintained for QTL analysis. QTL analysis was performed by Composite Interval Mapping, CIM (map function= kosambi, Model= 4.2) with a permutation time of 1000. QTL ICI Mapping was used for the construction of detailed linkage maps showing positions of QTLs. Due to the highly dense linkage maps generated, the construction of maps showing QTL positions focused on markers within QTL regions only. QTLs were named following nomenclature proposed by [48]. QTL names started with a lowercase “*q*”, which was followed by 2-3 initials of the corresponding measured trait (in upper cases) and then followed by the chromosome number. If there were more than one QTL on any chromosome, a “.” was used to uniquely identify them.

## Conclusion

This study revealed substantial genetic variation among forty-five F₂ cowpea populations derived from a cross between ‘Golinga’ and a wild relative using SNP markers, morphological traits, amino acids, and polyphenols. A total of seventy-nine QTLs were identified across four linkage groups, explaining up to 27.61% of the phenotypic variation. Seed-related QTLs consistently mapped to chromosome 4, whereas many vegetative, reproductive, and amino acid traits were concentrated on chromosome 1, highlighting these regions as key genomic hotspots.

Forty-two QTLs were associated with amino acids and nine with phenolic acids, demonstrating the polygenic nature of nutritional traits in cowpea. The detection of major QTLs for seed weight, and phytochemical composition provides valuable targets for marker-assisted selection. Overall, the SNP markers and QTLs identified in this study offer important baseline information for accelerating genetic improvement and advancing breeding strategies for cowpea.

## Acknowledgement

We are grateful for the support provided for this research by the Department of Plant and Environmental Biology, University of Ghana, Legon, Ghana Standard Authority and the Laboratory of the Government Chemist, United Kingdom.

## References

1. Begna T, Yesuf H. Genetic mapping in crop plants. Open Journal of Plant Science [Internet]. 2021;6(1):19–26. Available from: https://www.agriscigroup.us/articles/OJPS-6-128.php

2. Stancheva IK, Geneva M, Radka M, Donkova M, Hristozkova I, Perfanova M, et al. Nutritional value of cowpea grain grown under different moisture regimes with dual inoculation. Bulgarian Journal of Soil Science. 2016;1:112–21.

3. Prinyawiwatkul W, McWatters K, Beuchat L, Phillips R. Cowpea flour: A potential ingredient in food products. Crit Rev Food Sci Nutr. 1996;36:413–36.

4. Cai R, Hettiarachchy NS, Jalaluddin M. High-performance liquid chromatography determination of phenolic constituents in 17 varieties of cowpeas. J Agric Food Chem. 2003;51:1623–7.

5. Adjei-Fremah S, Jackai LE, Worku M. Analysis of phenolic content and antioxidant properties of selected cowpea varieties tested in bovine peripheral blood. Am J Anim Vet Sci. 2015;10(4):235–45.

6. Feng X, Yu X, Fu B, Wang X, Liu H, Pang M, et al. A high-resolution genetic linkage map and QTL fine mapping for growth-related traits and sex in the Yangtze River common carp (Cyprinus carpio haematopterus). BMC Genomics. 2018 Apr 2;19(1).

7. Mohammadi R, Mendioro MS, Diaz GQ, Gregorio GB, Singh RK. Genetic analysis of salt tolerance at seedling and reproductive stages in rice (O ryza sativa). Plant Breeding. 2014;133(5):548–59.

8. Slattery HD, Atwell BJ, Kuo J. Relationship between Colour, Phenolic Content and Impermeability in the Seed Coat of various Trifolium subterraneum L. genotypes. Ann Bot [Internet]. 1982 Sep 1;50(3):373–8. Available from: 10.1093/oxfordjournals.aob.a086376

9. Gupta PK, Rustgi S, Kumar N. Genetic and molecular basis of grain size and grain number and its relevance to grain productivity in higher plants. Genome. 2006;49:565–71.

10. Moura JDO, Rocha MDM, Lúcia R et al. Path analysis of iron and zinc contents and other traits in cowpea. Crop Breeding and Applied Biotechnology. 2012;12:245–52.

11. Burgess KS, Etterson JR, Galloway LF. Artificial selection shifts flowering phenology. Heredity (Edinb). 2007;99:641–8.

12. Greenup A, Peacock WJ, Dennis ES, Trevaskis B. The molecular biology of seasonal flowering-responses in Arabidopsis and cereals. Ann Bot. 2009;103(8):1165–72.

13. Orjuela J et al. A universal core genetic map for rice. Theoretical and Applied Genetics. 2010;120:563–72.

14. Ouédraogo JT et al. Development of SCAR markers linked to Striga race-specific resistance in cowpea. Afr J Biotechnol. 2012;11:12555–62.

15. Crepieux S, Lebreton C, Servin B, Charmet G. Quantitative Trait Loci detection in multi-cross inbred designs. Genetics. 2004;168:1737–49.

16. Crepieux S, Lebreton C, Flament P, Charmet G. Application of a new IBD-based QTL mapping method in wheat breeding. Theoretical and Applied Genetics. 2005;111:1409–19.

17. Muranty H. Power to test QTL detection using full-sib families under different schemes. Heredity (Edinb). 1996;76:156–65.

18. Moose SP, Mumm RH. Molecular plant breeding as the foundation for 21st century crop improvement. Plant Physiol. 2008;147:969–77.

19. Pan L, Wang N, Wu Z, Guan F, He M, Yang Z, et al. Genome-wide SNP identification and association analysis for yield-related traits in cowpea (Vigna unguiculata). BMC Genomics [Internet]. 2017;18:1–13. Available from: 10.1186/s12864-017-4121-6

20. Lo S, Muñoz-Amatriaín M, Boukar O, Herniter I, Cisse N, Guo YN, et al. Identification of QTLs for domestication-related traits in cowpea (Vigna unguiculata) using a high-density SNP-based linkage map. Theoretical and Applied Genetics [Internet]. 2018;131(12):2937–53. Available from: 10.1007/s00122-017-2999-8

21. Zeng ZB. Precision mapping of quantitative trait loci. Genetics [Internet]. 1994;136(4):1457–68. Available from: https://www.genetics.org/content/136/4/1457

22. Pottorff M et al. Leaf morphology QTLs and candidate genes in cowpea. BMC Genomics. 2012;13:234.

23. Huynh BL, Ehlers JD, Huang BE, Muñoz-Amatriaín M, Lonardi S, Santos JRP, et al. A multi-parent advanced generation intercross (MAGIC) population for genetic analysis and improvement of cowpea. Plant Journal. 2018;98:1129–42.

24. Muchero W, Roberts PA, Diop NN et al. Genetic architecture of delayed senescence, biomass, and grain yield under drought stress in cowpea. PLoS One. 2013;8:e70041.

25. Xu P et al. Development of microsatellite markers and phylogenetic analysis in asparagus bean. Molecular Breeding. 2010;25:675–84.

26. Andargie M, Pasquet RS, Gowda BS, Muluvi GM, Timko MP. Molecular mapping of QTLs for domestication-related traits in cowpea. Euphytica. 2014;200:401–12.

27. Xu P et al. QTL mapping and epistatic interaction analysis in asparagus bean. BMC Genet. 2013;14:4.

28. Herniter IA, Lo S, Muñoz-Amatriaín M, Dossa K, Fatokun C, Close TJ, et al. Morphological trait genetics in cowpea. Theoretical and Applied Genetics. 2019;132:207–23.

29. Ishiyaku MF, Singh BB, Craufurd PQ. Inheritance of time to flowering in cowpea (Vigna unguiculata (L.) Walp.). Euphytica. 2005;142(3):291–300.

30. Andargie M, Pasquet RS, Muluvi GM, Timko MP. Quantitative trait loci analysis of flowering time-related traits identified in recombinant inbred lines of cowpea. Genome. 2013;56(5):289–94.

31. Paudel D, Dareus R, Rosenwald J, Muñoz-Amatriaín M, Rios EF. Genome-wide association study reveals candidate genes for flowering time in cowpea. Front Genet. 2021;12:667038.

32. Mohammed SB, Ongom PO, Belko N, Umar ML, Muñoz-Amatriaín M, Huynh BL, et al. Identification of stable QTLs for flowering time and maturity across phosphorus environments in cowpea. Genes (Basel). 2025;16:1–34.

33. Wu F, Price BW, Haider W, Seufferheld G, Nelson R, Hanzawa Y. Functional and evolutionary characterization of CONSTANS gene family in soybean. PLoS One. 2014;9:e85754.

34. Zhang B, Feng M, Zhang J, Song Z. CONSTANS-like proteins in plant flowering and abiotic stress responses. Int J Mol Sci. 2023;24:16585.

35. Cheng XF, Wang ZY. Overexpression of COL9 delays flowering by reducing expression of CO and FT in Arabidopsis. Plant Journal. 2005;43:758–68.

36. Lin K, Zhao H, Gan S, Li G. ELF4-like proteins EFL1 and EFL3 influence flowering time. Gene. 2019;700:131–8.

37. Agbicodo EM, Fatokun CA, Muranaka S, Visser RGF, van der Linden CG. Breeding drought tolerant cowpea: constraints, accomplishments, and prospects. Euphytica. 2009;167:353–70.

38. Ehlers JD, Hall AE. Cowpea (Vigna unguiculata L. Walp.). Field Crops Res. 1997;53:187–204.

39. Chen L, Liu YG. Male sterility and fertility restoration in crops. Annu Rev Plant Biol. 2014;65:579–606.

40. Rasheed A, Rehman AU, Ogbonnaya FC, others. Genetic dissection of seed-set and fertility traits under heat and drought stress. Plant Biotechnol J. 2018;16(5):926–38.

41. Fatokun CA, Menancio-Hautea DI, Danesh D, Young ND. Evidence for orthologous seed weight genes in cowpea and mung bean based on RFLP mapping. Genetics. 1992;132:841–6.

42. Iyakaremye JP, Fields J. Quantitative Trait Loci (QTL) Mapping for Protein Content, and Amino Acids in a Cowpea MAGIC Population. 2025;

43. Kongjaimun A, Kaga A, Tomooka N, Somta P, Shimizu T, Shu Y, et al. An SSR-based linkage map of yardlong bean and QTL analysis of pod length. Genome. 2012;55(2):81–92.

44. IBPGR. Descriptors for Cowpea. Rome: IBPGR Secretariat; 1983.

45. Singleton VL, Orthofer R, Lamuela-Raventos RM. Analysis of total phenols and antioxidants using Folin–Ciocalteu reagent. Methods Enzymol. 1999;299:152.

46. Wang J, Li H, Zhang L, Meng L. Users’ Manual of QTL IciMapping. Beijing, China; Mexico City, Mexico; 2014.

47. Wang J, Li H, Zhang L, Meng L. QTL IciMapping User’s Manual. Beijing and Mexico; 2014.

48. McCouch SR, Chen X, Panaud O, Temnykh S, Xu Y, Cho YG et al. Microsatellite marker development, mapping and applications in rice genetics and breeding. Plant Mol Biol. 1997;35:89–99.

